# How flagella glycosylation of the phytopathogenic bacteria *Pseudomonas amygdali* pv. tabaci 6605 affect transport and deposition in saturated sandy porous media

**DOI:** 10.1101/2025.10.22.683916

**Authors:** Xin Zheng, Mounia Achak, Edvina Lamy, Yannick Rossez

## Abstract

To mitigate bacterial contamination in underground farmland, a comprehensive understanding of the transport and adhesion mechanisms of phytopathogenic bacteria in porous media is crucial for safeguarding soil and groundwater. This study aims to elucidate the effects of *Pseudomonas amygdali* pv. tabaci 6605 flagella (Wild type, Δ*fliC* strain) and its glycosylation (Δ*fgt1* and Δ*fgt2* strains) on bacterial transport and deposition in sandy porous media through a combination of experimental observations and numerical simulations. Flagella play a key role in bacterial transport and deposition dynamics through its surface properties. Its intrinsic hydrophobicity enhances bacterial adhesion and promotes deposition onto sandy grains, while simultaneously limiting transport through the porous medium. However, glycosylation of flagellin introduces hydrophilic glycans, which counteract this effect by increasing the overall hydrophilicity of the bacterial surface. As a result, glycosylated flagella facilitate bacterial mobility and improves recovery in the effluent, while reducing retention within the sand matrix. These findings highlight the critical influence of flagella’s biochemical modifications on bacterial behavior in porous environments. They provide valuable insights for understanding and managing microbial contamination in subsurface systems.

**Importance:** This work, conducted using homogeneous laboratory sand, could be extended to other types of abiotic media found in natural environments, such as clay, heterogeneous sands, and soils. Our study highlights the impact of flagellar glycosylation on bacterial behavior, an essential factor for assessing the risk posed by phytopathogenic bacteria in agricultural settings and for developing effective soil bioremediation strategies. Moreover, this study provided valuable insights into the mechanisms governing bacterial transport and deposition at the macroscopic (column) scale under dynamic flow conditions. Investigating unsaturated flow conditions, which better approximate real field scenarios, may further our understanding of bacterial interactions at air/solid/water interfaces. Future research should explore bacterial movement across different spatial scales. In particular, pore-scale experiments can provide direct evidence of processes such as attachment and motility. This could significantly enhance our understanding of microbial dynamics in complex environments.

## Introduction

The dissemination of phytopathogenic bacteria through subsurface environments such as soil and water is a critical step in plant infection but remains poorly understood. Numerous studies have investigated how environmental and bacterial factors influence transport and retention in porous media. These include physical parameters of the medium (1–3), fluid properties (4–6), and cellular characteristics such as morphology, size, and surface charge (7–10). Among these, flagella-driven motility has emerged as a pivotal attribute influencing bacterial transport and attachment.

Bacterial flagella are complex structures composed of a rotary motor, hook, and a long extracellular filament that functions as a helical propeller (11). In addition to driving motility, flagellar filaments play a critical role in surface attachment, often serving as the first point of contact between the cell and a surface (12, 13). In many bacterial species, the filament is primarily composed of a protein called flagellin, encoded by the *fliC* gene. In *Pseudomonas syringae*, a known plant pathogen, flagellar motility and the expression of *fliC* are essential for virulence and efficient colonization (14–16). Beyond gene presence and expression, post-translational modifications (PTMs) of flagellin, particularly glycosylation and methylation, have been shown to modulate bacterial motility, adhesion, and host interactions. For instance, in *Salmonella enterica* serovar Typhimurium, methylation of lysine residues on flagellins enhances transport and adhesion to sandy surfaces under flow conditions(17). This finding raises the question of whether flagella glycosylation, another widespread PTM, similarly influences bacterial behavior in hydrodynamic environments. Flagellin glycosylation, the covalent addition of sugars to amino acid residues, has been identified in numerous bacterial species and can affect protein stability, folding, immune recognition, and function (18–20). However, its role in bacterial transport through abiotic porous media under dynamic conditions remains largely unexplored.

To address this knowledge gap, we focused on *Pseudomonas amygdali* pv. tabaci *6605,* formerly classified as *P. syringae* pv. tabaci (21), a phytopathogen responsible for wildfire disease in tobacco. This organism produces a single flagellin type (FliC), which is O-glycosylated with rhamnose-rich glycans and a terminal modified 4-amino-4,6 dideoxyglucosyl (Qui4N) residue,β-D-Qui*p*4N(3-hydroxy-1-oxobutyl)2Me, commonly referred to as viosamine (mVio) (22). The biosynthesis of this glycan structure is regulated by two genes*, fgt1* and *fgt2*; deletion mutants of these genes produce non-glycosylated or partially glycosylated flagellin, respectively (23). Previous work has shown that loss of flagella glycosylation in this strain reduces swarming motility and virulence (23, 24), suggesting a potential role in modulating surface interactions.

While protein glycosylation has been linked to altered bacterial adhesion to biotic surfaces (25–27), little is known about its role in bacterial transport and deposition onto abiotic surfaces under hydrodynamic conditions. This study aims to address that gap by assessing whether flagella glycosylation affects bacterial transport and retention in porous media under flow. In this study, we examined how the presence and glycosylation state of flagella affect bacterial transport and retention in saturated flow through homogeneous sandy porous media. Using *Pseudomonas amygdali* pv. tabaci 6605 as a model, we compared the wild-type strain with three isogenic mutants: Δ*fliC*, lacking flagellin and flagella; Δ*fgt1*, which produces non-glycosylated flagella; and Δ*fgt2*, which expresses partially glycosylated flagella. Column-scale transport experiments were conducted under controlled laboratory conditions, and breakthrough curves (BTCs) were generated to assess bacterial movement through the porous medium. These BTCs were analyzed using the Hydrus-1D code with a mobile–immobile (MIM) model to evaluate how differences in flagella structure and modification affect bacterial transport and filtration dynamics. In addition to genetic and structural differences, we also investigated the role of flagellar surface hydrophobicity, a property influenced by glycan composition, on bacterial deposition. By integrating experimental observations with hydrodynamic modeling, this study provides new insights into the contribution of flagella and its glycosylation to bacterial motility, adhesion, and environmental dissemination in dynamic soil and water systems.

## Materials and methods

### Bacterial strains

The *P. amygdali* pv. tabaci strains used in this study (Table 1) include the wild type, a Δ*fliC* mutant lacking flagella, and two glycosyltransferase mutants: Δ*fgt1*, which produces non-glycosylated flagella, and Δ*fgt2*, which expresses partially glycosylated flagella. In the wild type, the flagellin polymer is O-glycosylated with glycans composed of two or three rhamnose residues and a terminal mVio, attached to six serine residues (positions 143, 164, 176, 183, 193, and 201) (22).

**Table 1.**
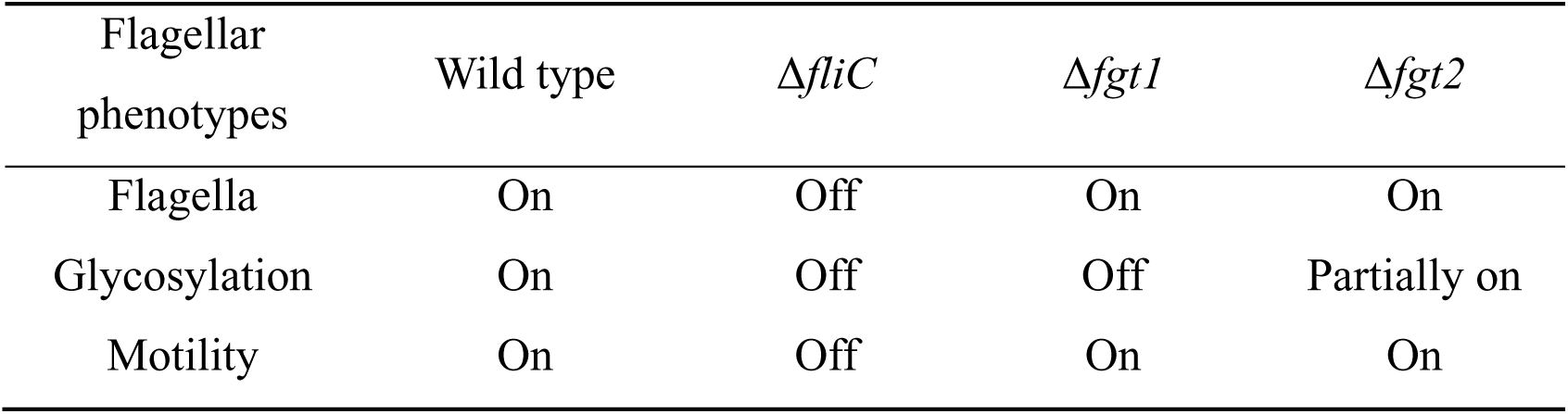
Cell properties for *P. amygdali* pv. tabaci strains.

### Bacterial suspension preparation

*P. amygdali* pv. tabaci strains were inoculated on a King’s medium B base broth (KB) agar at 28 °C. Then, these strains were grown at 28 °C, 85 rpm in KB. 20 g proteose peptone, 1.5 g K_2_HPO_4_, 15 mL glycerol and 1.5 g MgSO_4_·7H_2_O were needed for preparing 1 L of KB, 20 g agar were required additionally to make KB agar. 1.5 g MgSO_4_·7H_2_O were added after autoclaving. Bacterial suspension harvested from the stationary phase of bacterial culture was used in this work to avoid bacterial growth during experiments. The numbers of initial and final bacteria were determined by the number of colony-forming units (CFU) using the plating method on KB agar as described previously (17).

### Flagella purification and surface hydrophobicity measurement

The flagella were purified (12, 28) and the surface hydrophobicity were measured as described previously (29). In short, *P. amygdali* pv. tabaci strains were cultured in KB media at 28℃ for 36 h in an orbital shaker incubator at 80 rpm, harvested by centrifugation then sheared off using a magnetic stirrer for 1h, followed by centrifugation to collect the supernatant containing the flagella. Ammonium sulfate was slowly added with vigorous stirring until two-thirds saturation was achieved. The flagella were harvested by centrifugation at 15000 *g* for 20 min under 4℃ after 36 h incubation, after that, the precipitation was re-suspended in Tris-buffered saline. The surface hydrophobicity (So) of purified flagella using PRODAN (30). 8 µL of 1 mM of PRODAN (prepared in DMSO) solution was added to 1 mL flagella samples which diluted in 20 mM HEPES and 150 mM NaCl (pH=7.4). The samples were incubated for 10 min in the dark and the fluorescence intensity was measured with a Cary Eclipse spectrofluorometer. The wavelengths of excitation and emission were 365 nm and 465 nm respectively, with corresponding 5 nm and 5 nm slits widths. The surface hydrophobicity values were determined using at least four measures per sample repeated three times and the mean value was used.

### Column experiments and numerical simulations

A Plexiglass column with an inner diameter of 3.4 cm and a length of 18 cm was used for the transport experiments. Homogeneous Fontainebleau sand, with particle sizes ranging from 0.16 mm to 0.79 mm (D_50_ = 0.36 mm), was used as the porous medium in this study. The bulk density, porosity, and degree of saturation of the columns were determined by gravimetric analysis (see Supplementary Material 1). A 20 mL bacterial suspension was injected into the columns, and breakthrough curves (BTCs) were generated by plotting the time-dependent bacterial concentration in the effluent. Moment analysis was used for estimating retardation factor (*R*) and mass balance from effluent (M_eff_) (31).

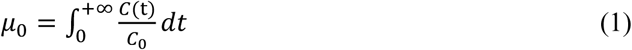

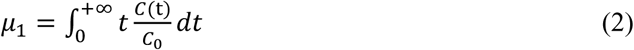

Here μ_0_ is zero-order moment of BTCs, μ_1_ is the first-order moment of BTCs, *C*(t) is the time-dependent concentration of the effluent, *C*_0_ is the initial concentration of bacteria. M_eff_ is calculated by the ratio of the bacteria mass recovered at the column outlet to their mass injected at the column inlet (32),

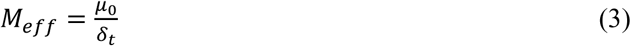

here δ_*t*_ is the time for injection bacteria suspension into the column. *R* was determined by the ratio of resident time for bacteria to the theoretical solution resident time,

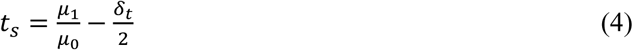

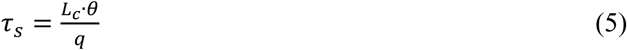

here *L*_*c*_ is the column length (cm), θ is the total water content, *q* is the Darcy velocity (cm/min).

After the transport experiments, the porous medium inside the column was sectioned into eight successive layers of 2.25 cm to analyze the spatial distribution of bacteria. Each sandy layer was transferred into a flask containing 0.1 mmol/L NaCl solution and shaken on a rotary shaker to detach bacteria from the sand grains. The plating method was used to quantify the bacterial biomass recovered from the retention profiles, denoted as M_retained_. The total recovery of bacteria (M_total_) is the sum of M_eff_ and M_retained_.

The modified two region mobile-immobile (MIM) model, incorporating two kinetic deposition sites as previously described (2, 17), was employed in this study. In the MIM model, the liquid phase is conceptualized as two distinct regions: a mobile (flowing) region and an immobile (stagnant) region. The model assumes that bacteria are excluded from the immobile regions and that no exchange occurs between the mobile and immobile zones. Column transport experiments in this study were conducted under saturated water conditions. Under these conditions, the two deposition sites were interpreted as bacterial attachment to solid–water interfaces (SWI) and irreversible straining, both considered key mechanisms of bacterial retention in porous media (Fig. 1) (33). A reversible detachment process was also considered. The mass transfer between the liquid and solid phases is described by the equation 6 (34):

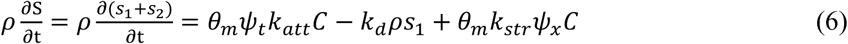

**Fig. 1.**
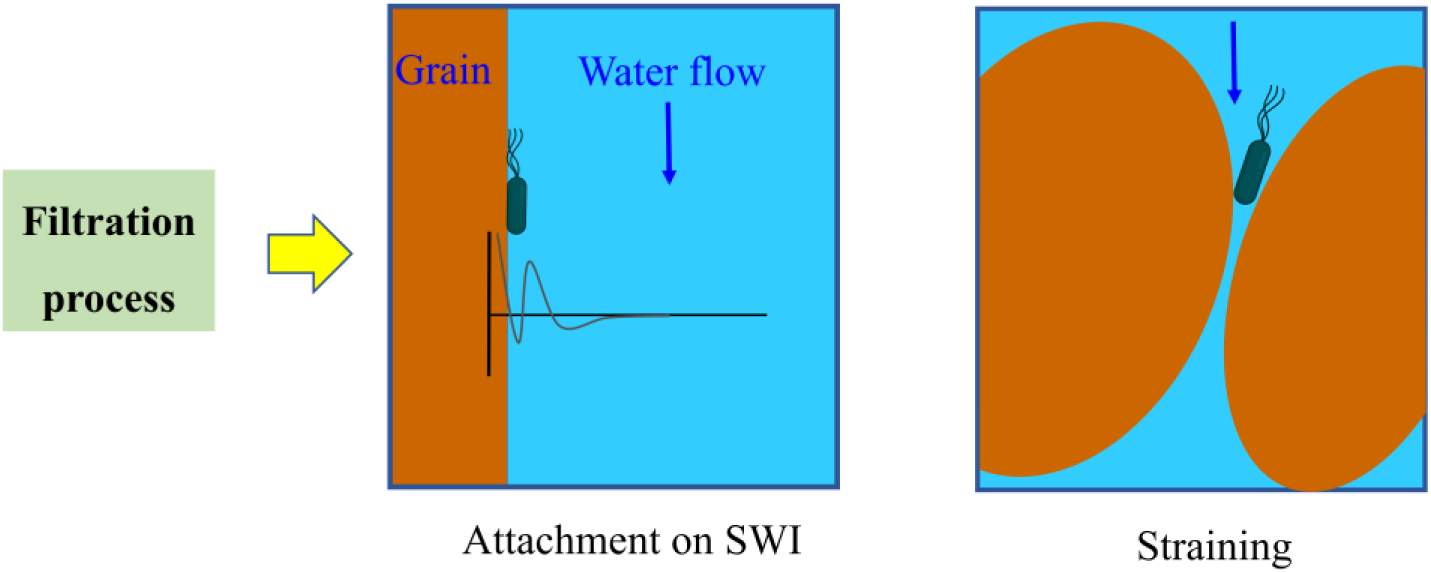
Mechanisms of bacterial deposition in columns under saturated conditions include two primary patterns: physicochemical attachment to solid-water interfaces (SWI), which allows for potential reversible detachment, and physical straining, which leads to irreversible retention.

Here *ρ* is the bulk density of the porous media (g/cm^3^), *S* is the bacterial concentration in solid phase (CFU/g), *s_1_* and *s_2_*are the bacterial concentrations in solid phase accounting for attachment or irreversible straining (CFU/g), respectively; *θ_m_* is the mobile water content, *C* is the bacterial concentration in the liquid phase (CFU/cm^3^), *t* is time (min), *x* is the distance (L); *k_att_* is the first-order attachment coefficient (1/min), *k_d_*is the first-order detachment coefficient (1/min), *k_st_*_r_ is the first-order straining coefficient (1/min); *Ψ_t_* and *Ψ_x_* describe the deposition of bacteria under time– and depth-dependent deposition, respectively.

### Statistical analysis

The statistics were analyzed for significance using Student’s *t-*test by SPSS 26.0. Origin 8.1 was used for graphic drawing.

## Results

### Flagellin expression reduces bacterial mobility and promotes retention within porous media

*P. amygdali* pv. tabaci WT (with flagella) and Δ*fliC* strains (without flagella) were used in this study to investigate the effects of flagella on bacterial transport and deposition in a sandy medium. Column experiments were conducted in triplicate and average BTCs (Fig. 2a) as well as retention profiles (RPs, Fig. 2b) displayed a good repeatability for each strain, the experimental parameters are listed in Table 2. Experimental conditions for all replicates are shown in Supplementary Material 2.

**Fig. 2.**
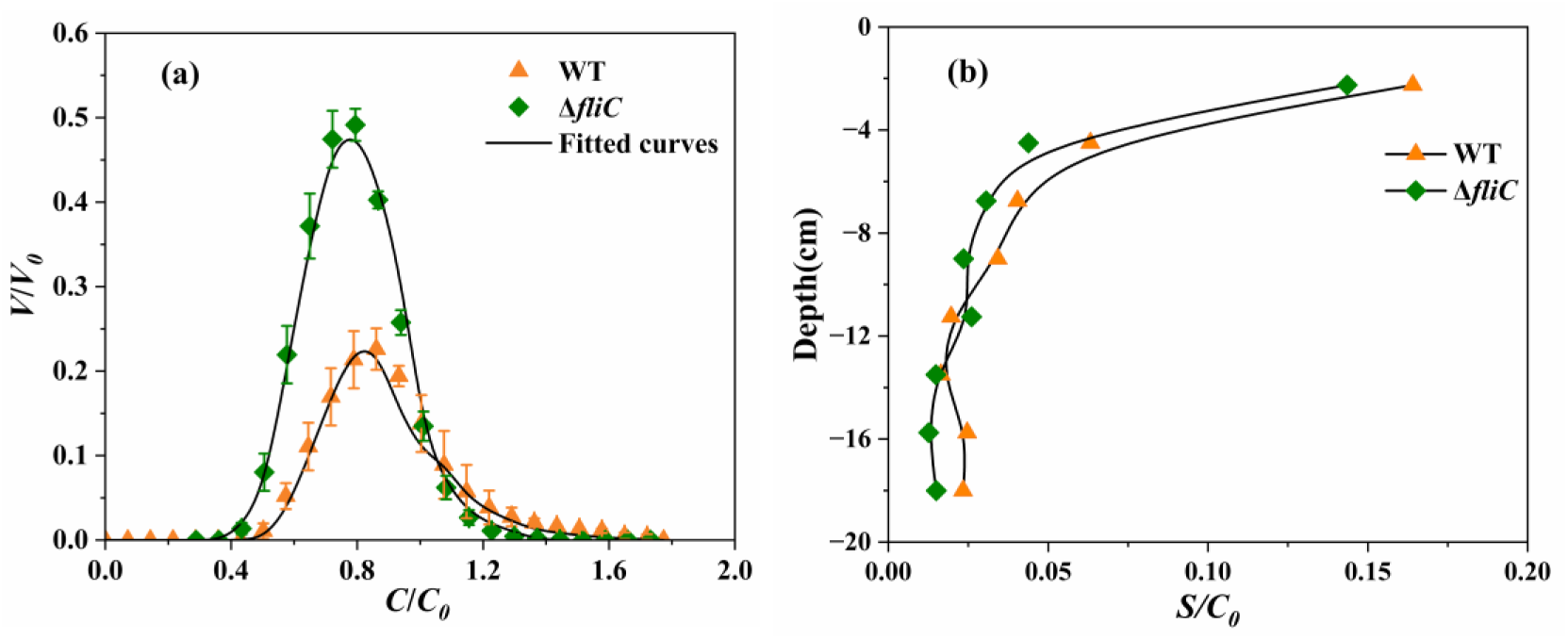
Observed (symbols) and fitted (curves) BTCs. (a) and RPs (b) of *P. amygdali* pv. tabaci WT (with flagella) and Δ*fliC* (without flagella) strains. *V* is the volume occupied by water in the column, *V*_0_ is the total pore volume of the column, and *C*, *C*_0_ and *S* are the time-dependent, initial and solid phase concentration of bacteria, respectively.

**Table 2.** Experimental parameters for P. *amygdali* pv. tabaci transport experiments conducted under saturated fluid flow conditions. Mass balance includes bacteria measured in the effluent (M_eff_) and bacterial biomass recovered from retention profiles (M_retained_), with total bacterial recovery (M_total_)

| Bacterial strains | $R$ | Recovery rate (%) | | |
| --- | --- | --- | --- | --- |
| | | $M_{eff}$ | $M_{retained}$ | $M_{total}$ |
| WT<br>(with flagella) | $0.70 \pm 0.06^a$ | $14.68 \pm 1.95$ | $65.04 \pm 1.01$ | $79.72 \pm 1.36$ |
| $\Delta fliC$<br>(without flagella) | $0.63 \pm 0.01$ | $29.24 \pm 1.32$ | $52.53 \pm 0.85$ | $81.77 \pm 2.15$ |
|  |  | ** | *** | NS |
| $\Delta fgt1$<br>(flagella without glycosylation) | $0.64 \pm 0.05$ | $10.56 \pm 1.07$ | $73.46 \pm 5.60$ | $84.03 \pm 4.81$ |
|  |  | NS | NS | NS |
| $\Delta fgt2$<br>(flagella partially glycosylated) | $0.69 \pm 0.01$ | $21.08 \pm 0.70$ | $59.45 \pm 3.52$ | $80.53 \pm 4.12$ |
|  |  | * | NS | NS |

<sup>a</sup>Values indicate standard deviations, with statistical significance determined by Student's *t*-test relative to the WT (\*\*\* = $p < 0.001$ ; \*\* = $p < 0.01$ ; \* = $p < 0.05$ ; NS = not significant) with $n=3$ .

Both WT and Δ*fliC* strains showed a symmetrical shape on BTCs (Fig. 2a), although the BTCs of Δ*fliC* strain exhibited a more symmetrical shape compared to the BTCs of WT. The peak of BTCs for both strains occurred before 1*V*/*V_0_*(pore volume) indicating a preferential transport of bacteria.

The Δ*fliC* strain exhibited faster transport than the WT strain, as indicated by its lower retardation factor (0.63 compared to 0.70 for WT; Table 2) and earlier breakthrough times. Different recovery rates from effluent and retention profiles were obtained for WT and Δ*fliC* mutants (Table 2). The average recovery of effluent for WT was 14.68% and 29.24% for Δ*fliC* strains, in agreement with the BTCs behavior, with a higher peak for Δ*fliC* strains. Logically, WT was more retained in the sand (65.04%) than Δ*fliC* strain (52.53%). Similar total recovery (effluent + profiles) was obtained for both strains, with lower total recovery for WT strain (79.72%), while Δ*fliC* strain reached 81.77%. The RPs replicates of WT and Δ*fliC* strains (Figure 2b) displayed a non-monotonic distribution behavior with a high number of attached bacteria at the column inlet (layer 0∼2.25cm). WT exhibited higher dispersion in deeper layers (12∼18 cm) than Δ*fliC* strains.

### Glycosylation of flagella promotes bacterial transport through the sand, resulting in higher effluent recovery and decreased retention

To explore the influence of flagella glycosylation on bacterial transport and deposition, Δ*fgt1* (without glycosylation) and Δ*fgt2* (partially glycosylated) strains were observed and compared in regard to BTCs and RPs, together with WT. BTCs and RPs of all strains showed a good repeatability (Fig. 3).

**Fig. 3.**
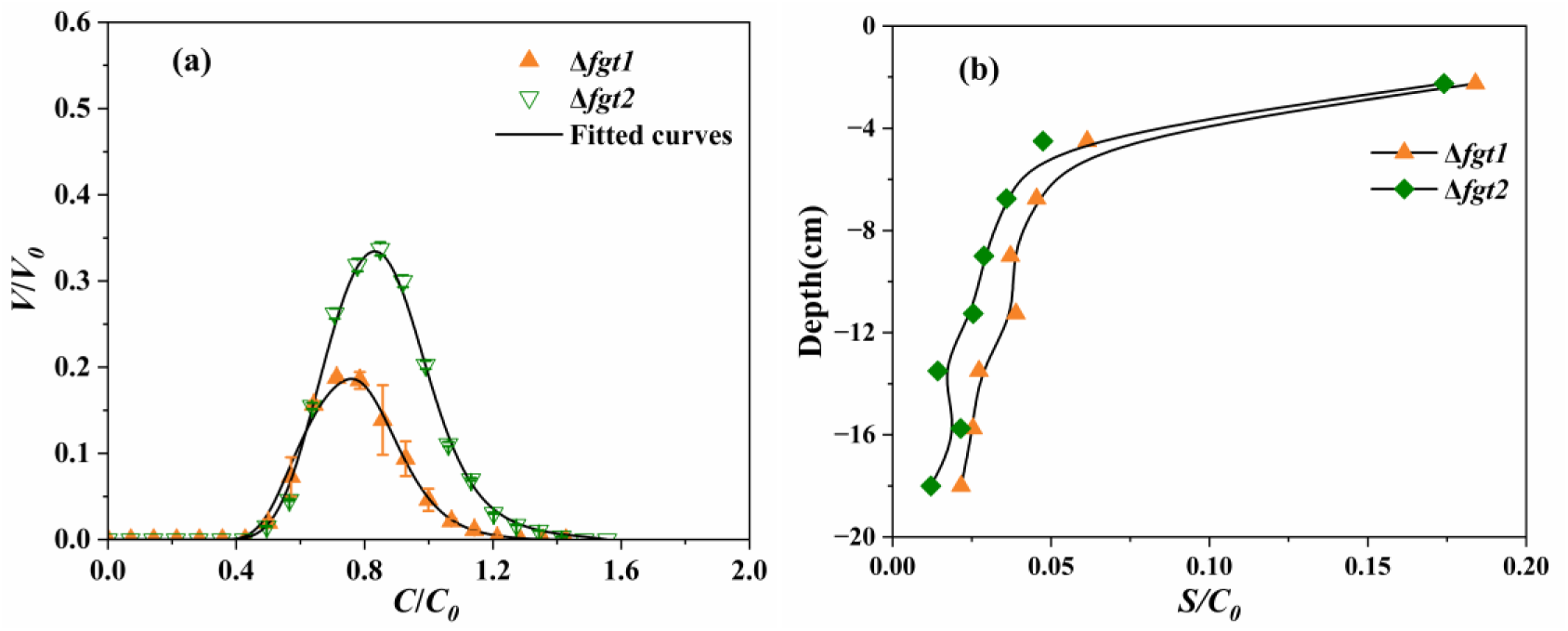
Observed (symbols) and fitted (curves) BTCs. (a) and RPs (b) of *P. amygdali* pv. tabaci Δ*fgt1* (flagella without glycosylation) and Δ*fgt2* (flagella partially glycosylated) strains.

All replicates showed a symmetrical shape on BTCs (Fig. 3a). The Δ*fgt1* strain lacking flagella glycosylation exhibited a faster transport than the WT and Δ*fgt2* (partially glycosylated) strain. This was confirmed by the same start of BTCs, but earlier end of BTCs. The same result was also illustrated by a lower retardation factor of Δ*fgt1* strain (0.64, Table 2) compared to WT (0.7) and Δ*fgt2* strain (0.69). Although no obvious difference was observed for the BTCs between WT and Δ*fgt2* strain, which displayed almost the same starts and ends of BTCs, the peak of Δ*fgt2* strain was higher than WT, suggesting higher recovery of this strain in the effluent, confirmed by higher M_eff_ (Table 2).

Even though the Δ*fgt1* exhibited a faster transport, this strain had the lowest recovery in the effluent (10.56%, Table 2) in comparison of WT (14.68%) and Δ*fgt2* strain (21.08%), a result in agreement with the lowest BTCs peak of replicates. The Δ*fgt1* strain was mostly retained in the sand (73.46%) compared to the WT and Δ*fgt2* strains presenting 65.04% and 59.49% retention rates respectively. Furthermore, Δ*fgt1* strain showed the highest total recovery (84.03%) compared to WT (79.72%) and Δ*fgt2* strain (80.53%).

The RPs of WT (Fig. 2b), Δ*fgt1* and Δ*fgt2* (Fig. 3b) strains had a non-monotonic distribution among replicates with highest number of bacteria retained (*S/C_0_*) in the column inlet (0∼2.25cm). The dispersion of Δ*fgt2* strain was the lowest among layers 2.25∼18 cm, which showed a better repeatability than WT and Δ*fgt1* strains. Δ*fgt1* strain was more dispersed among replicates in layer 2.25∼18 cm. Contrary to Δ*fgt1* and Δ*fgt2* strains, WT had a more dispersed number of bacteria in layer 12∼18 cm.

### Flagella hydrophobicity reduces bacterial transport but promotes deposition

*P. amygdali* pv. tabaci WT, Δ*fgt1* and Δ*fgt2* were observed to explore the effects of flagella hydrophobicity on bacterial recovery. The surface hydrophobicity of the purified flagella from WT, Δ*fgt1* and Δ*fgt2* (Fig. 4a), was higher for Δ*fgt1* mutant than that of WT and showed a significant difference (P < 0.001). No significant difference was observed between WT and Δ*fgt*2 strains.

**Fig. 4.**
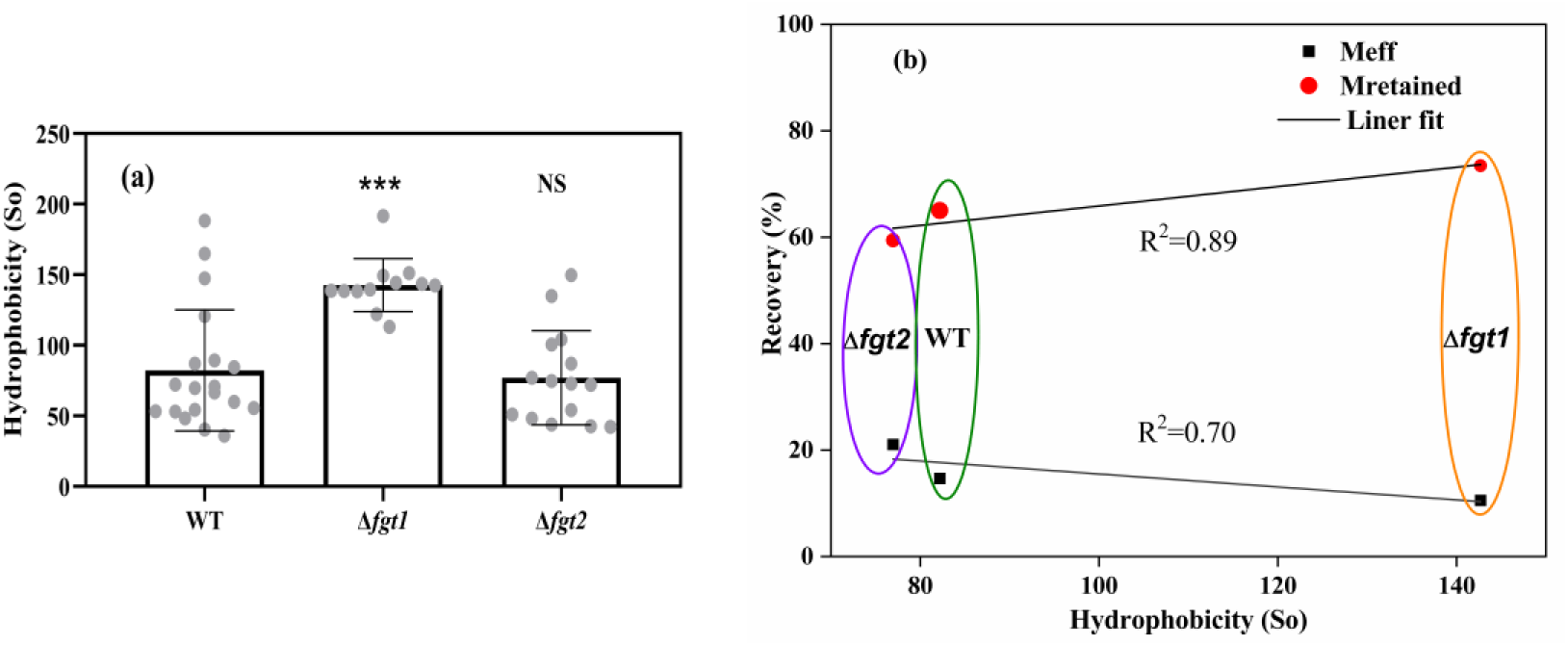
(a) The flagella hydrophobicity of *P. amygdali* pv. tabaci WT, Δ*fgt1* (without glycosylation) and Δ*fgt2* (partially glycosylated) strains. Replicates are shown as mean values, error bars represent standard deviations, and statistical significances were determined by the student’s t-test with WT (*** = *p*< 0.001; NS = not significant); (b) Relationship between bacterial flagella hydrophobicity, recovery rate from effluent (M_eff_) and retention rate in the sand (M_retained_).

A linear relationship was observed between flagella hydrophobicity and both the effluent recovery rate (Meff) and retention rate in sand (Mretained) (Fig. 4b). Flagella hydrophobicity showed a strong positive correlation with bacterial retention in the sand (*R²* = 0.89), indicating that increased hydrophobicity enhances adhesion to the sand matrix. Conversely, a moderate negative correlation was observed between hydrophobicity and effluent recovery (*R²* = 0.70), suggesting that more hydrophobic strains are less efficiently transported through the porous medium. Among the three flagellated strains, the Δ*fgt1* mutant, characterized by the highest flagella hydrophobicity, exhibited the lowest recovery in the effluent and the highest retention in the sand compared to the wild type and Δ*fgt2*. These results suggest that flagella hydrophobicity is a key factor driving bacterial deposition in porous media.

### Bacteria preferential transport and their mechanisms of deposition

The BTCs for all mutants of *P. amygdali* pv. tabaci were well described using the MIM model based on Hydrus-1D code (Fig. 2 and Fig. 3) with R^2^>0.93. Mean values of transport and deposition parameters are reported in Fig. 5 and the fitted parameters of all replicates are shown in Supplementary Material 3.

**Fig. 5.**
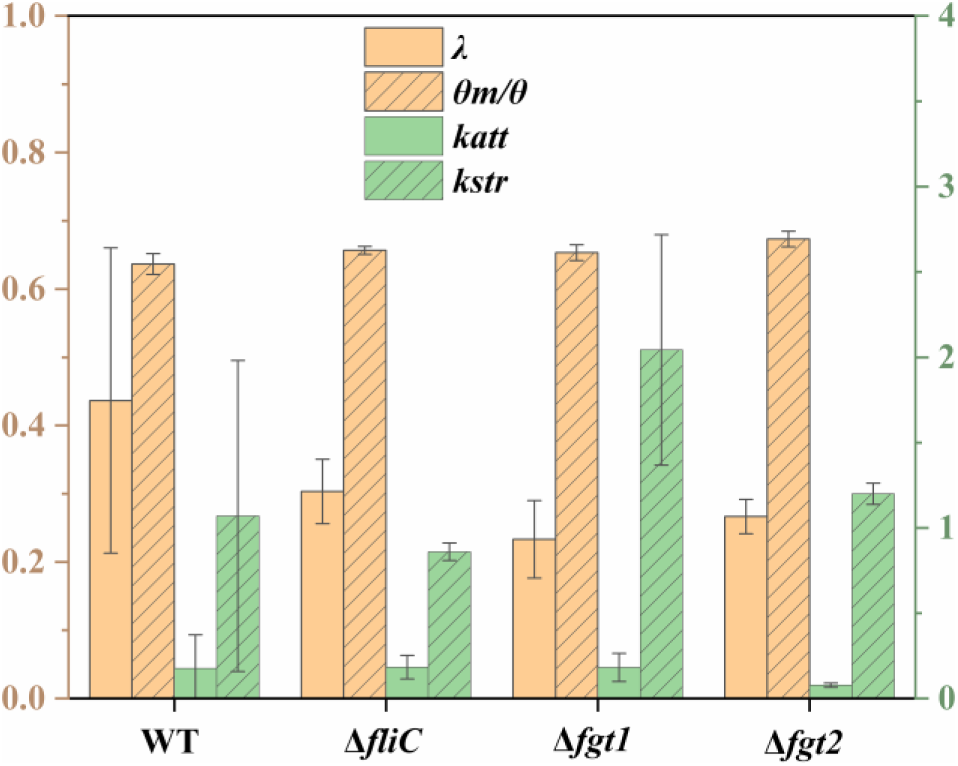
Fitted parameters of *P. amygdali* pv. tabaci WT, Δ*fliC*, Δ*fgt1* and Δ*fgt2* strains. Replicates are shown as mean values, error bars represent standard deviations.

The dispersivity *λ* and the mobile water fraction *θ*_m_/*θ* are key indicators of flow and transport behavior in porous media. Low *λ* and high *θ*_m_/*θ* values suggest more uniform, less dispersive flow with reduced preferential transport pathways. All bacterial strains showed similar transport behavior, with mean *θ*_m_*/θ* values, ranging from 0.64 to 0.68 (Fig. 5), indicating that 64–68% of the pore volume was involved in convective transport, while the remaining 32–36% represented immobile zones that were either excluded from flow or available only for diffusive transport. No significant differences in *θ*_m_*/θ* were observed among the strains. Dispersivity *λ* was highest for the wild-type strain (0.44 cm), followed by Δ*fliC* (0.33 cm), Δ*fgt2* (0.27 cm), and Δ*fgt1* (0.23 cm). Although the WT strain showed greater variability between replicates, all λ values remained within the same order of magnitude, suggesting broadly comparable transport dynamics across strains.

The first-order attachment (*k_att_*), detachment (*k_d_*) and straining (*k_str_*) coefficients derived from simulations highlight the respective contributions of physicochemical attachment, reversible detachment, and physical straining to bacterial deposition. The WT, Δ*fliC*, and Δ*fgt1* strains exhibited similar attachment coefficients, with a mean *k_att_*value of 0.18 min⁻¹, indicating comparable attachment behavior. In contrast, the Δ*fgt2* mutant displayed a significantly reduced *k_att_* value (0.08 min^-1^) (Fig. 5, Supplementary Material 3), approximately one order of magnitude lower, suggesting impaired attachment efficiency likely due to partial glycosylation of flagella. The WT strain exhibited a significantly lower detachment coefficient *k_d_* 0.39 min^-1^ compared to the Δ*fliC* mutant, which showed a value of 3.06 min⁻¹ (Supplementary Material 3). This indicates that the Δ*fliC* strain is more prone to reversible detachment from sandy grains under hydrodynamic forces. A similar trend was observed for the Δ*fgt1* and Δ*fgt2* mutants with *k_d_*of 2.05 and 1.29 min^-1^ respectively. These findings suggest that the WT strain is more likely to undergo irreversible attachment, likely driven by stronger physicochemical interactions with the sand surface. The *k_str_* values obtained for all bacterial strains ranged from 0.86 to 2.04 min^-1^ (Fig. 5). Given the similar orders of magnitude and standard deviations of the *k*_str_ values across all strains, physical straining appears to be a consistent mechanism contributing equally to bacterial retention among the tested strains. As previously reported by Zheng et al.(17), some parameters derived from numerical simulations exhibited standard error coefficients (S.E. Coeff., obtained from the HYDRUS-1D model; see Supplementary Material 3) that were comparable in magnitude to their corresponding absolute values. Nevertheless, comparisons between the different mutants can still reveal qualitative trends that may be linked to their distinct biological characteristics.

## Discussion

Flagella not only drives flagellar motility but also functions as a factor in bacterial adhesion to host cells (13, 15, 35, 36) and other surfaces, including polystyrene (37). In this study, the Δ*fliC* strain lacking flagella was the most efficiently recovered in the effluent and exhibited the lowest retention within the sand (Table 2). Lower retardation factors and earlier breakthroughs observed for the Δ*fliC* strain relative to the wild type (Fig. 2a, b) suggest faster transport, potentially due to the absence of flagella and the resulting loss of motility. The Δ*fliC* mutant exhibits a significantly reduced capacity to elicit disease symptoms compared to the wild-type strain (38). In agreement with the present work, the *S*. Typhimurium Δ*flgKL* mutant, lacking flagella, demonstrated faster transport through porous media than both the WT and the non-motile, flagellated Δ*motAB* strain (17). This mutant also showed lower retention in sand compared to flagellated strains, further supporting the role of flagella in promoting surface interactions. Similarly, previous studies have reported that non-flagellated bacterial strains exhibit greater mobility than flagellated ones in sandy media (39). Overall, the Δ*fliC* mutant in our study exhibited greater mobility, likely due to reduced surface adhesion, resulting in shorter residence times and more efficient transport through porous media.

The WT strain showed more dispersive flow behavior, as indicated by a higher mean dispersivity relative to the Δ*fliC* strain, which may be attributed to the influence of flagella on bacterial movement and interaction with the porous medium (17). Lower mean dispersivity and more uniform flow patterns, reflected by higher *θ*_m_/*θ* values, were observed for the *S. Typhimurium* Δ*flgKL* mutant lacking flagella, compared to both the wild-type and the non-motile, flagellated Δ*motAB* strain in porous media. These differences were attributed to the presence or absence of flagella, suggesting that flagellated bacterial strains tend to exhibit less uniform flow and more preferential transport pathways in porous environments. While a similar trend in dispersivity was observed in our study, the close *θ*_m_/*θ* values between the WT (64%) and Δ*fliC* (66%) suggest only minor differences in flow uniformity, making it difficult to establish a clear link between flagella presence and preferential transport behavior. Post-translational modification of flagella significantly influenced bacterial transport, as reflected by effluent recovery rates. The Δ*fgt1* strain, lacking flagella glycosylation, exhibited the lowest recovery in the effluent compared to the glycosylated wild-type and the partially glycosylated Δ*fgt2* strain. Glycosylation may enhance the structural stability of the flagellar filament by stabilizing polymerized flagellin proteins, as reported for *P. amygdali* pv. *tabaci*(*24*). This structural reinforcement could explain the reduced recovery and increased retention of the non-glycosylated strain in the sand matrix. As observed for flagella presence, its post-translational modification via glycosylation did not appear to influence flow uniformity under the experimental conditions of this study, as indicated by the similar *θ*_m_/*θ* values across strains. The *P. amygdali* pv. tabaci ΔfliC mutant exhibited the lowest deposition in porous media among all tested strains, likely due to the absence of flagella, which may reduce its adhesion capacity in the sand matrix. This observation is consistent with previous findings showing that *E. coli* O157:H7 strains lacking FliC exhibited significantly lower adhesion to bovine intestinal and plant tissues compared to the wild type (12, 40). Moreover, non-flagellated *S*. Typhimurium and *E. coli* mutants exhibited reduced deposition onto sand grains compared to the WT strain (17, 41). Our findings underscore the significance of flagella in facilitating bacterial deposition by augmenting surface adhesion, both *in vitro* and *in vivo*. The mechanisms driving adhesion of the WT strain to surfaces remain to be fully elucidated. In many motile bacteria, adhesion often involves the formation of non-motile, multicellular communities called biofilms, which enable stable surface attachment and enhanced environmental persistence (42). In these biofilms, adhesins such as pili and flagella, as well as extracellular matrix components like DNA and exopolysaccharides, along with bacterial motility, play crucial roles in early attachment and biofilm development. This presents a greater challenge for bacteria to detach from surfaces (43–45), thereby enabling them to thrive and persist in diverse environments (46).

The WT strain exhibited a lower retention rate compared to the non-glycosylated Δ*fgt1* mutant, highlighting the critical role of flagella glycosylation in enhancing bacterial adhesion and biofilm formation (27, 47). The swimming activity of glycosylation-defective mutants was prominently decreased in a highly viscous medium (48). However, the motility of glycosylation-defective mutants was not affected compared to WT strain (Supplementary Material 4). *P. amygdali* pv. tabaci strains deficient in flagella glycosylation showed diminished swarming motility and impaired biofilm formation on tobacco plant surfaces. Additionally, their adhesion to polystyrene surfaces was significantly reduced (23). Swarming, a collective form of surface motility, allows bacterial populations to rapidly migrate across solid substrates, facilitating colonization and the establishment of dense communities (49–51). Given the well-established links between bacterial motility (including swimming and swarming), surface adhesion, and biofilm formation, the observed reduction in both adhesion and swarming motility in the non-glycosylated strain likely impairs its ability to initiate and sustain biofilms within porous media during column experiments. This reduced biofilm-forming capacity may underlie the altered transport dynamics and lower retention observed for this strain. The interplay between glycosylation, motility, and surface attachment underscores the pivotal role of glycan modifications in shaping bacterial interactions with complex environments. These findings highlight glycosylation not merely as a structural trait, but as a key functional factor influencing bacterial colonization, persistence, and spatial organization in heterogeneous systems such as porous substrates. The non-glycosylated Δ*fgt1* strain exhibited a higher propensity for detachment from sandy grains, which may explain its elevated detachment coefficient (*k_d_*) compared to the glycosylated WT and Δ*fgt2* strains. The Δ*fgt1* strain demonstrated greater retention than the WT strain due to physicochemical attachment forces, a process that is reversible. Additionally, the Δ*fgt1* strain could still be retained through physical straining, as evidenced by the higher *k_str_*values observed in our study compared to the other strains (Fig. 5). Our findings reveal that bacterial strains deficient in flagella glycosylation display the highest surface hydrophobicity (Fig. 4a), This observation aligns with previous studies demonstrating that the presence of pilin glycans in *Pseudomonas aeruginosa* reduces the hydrophobicity of both purified pili and whole cells (52). As anticipated, the observed positive correlation between flagella hydrophobicity and retention rates (Fig. 4b) aligns with previous research findings on the relationship between bacterial surface hydrophobicity and retention under unsaturated flow conditions (53). Hydrophobicity was identified as the primary factor influencing bacterial deposition. The Δ*fgt1* strain demonstrated higher hydrophobicity compared to the WT, resulting in increased deposition on sandy surfaces. In contrast, the Δ*fgt2* strain exhibited flagellar hydrophobicity like that of the WT, which led to enhanced deposition relative to the WT strain. Similarly, *S*. Typhimurium lacking flagella methylation showed significantly reduced deposition compared to the WT (17), with its flagellar hydrophobicity being lower than that of the WT (29). Research has extensively explored flagellar post-translational modifications, particularly in bacteria from natural aquatic environments such as *Shewanella oneidensis* (54) and various other bacterial species including Enterobacteriaceae (20, 55). This study underscores the critical role of flagellar glycosylation in promoting the dissemination of plant pathogens via water runoff and their potential transport into groundwater systems. These post-translational modifications may enhance bacterial motility and surface interactions, offering an advantage in colonizing diverse environments by finely tuning the balance between dispersal and biofilm formation. However, our understanding of how glycosylation contributes to the environmental dissemination of pathogens remains limited. A key challenge lies in dissecting the specific contributions of glycosylation, as current approaches such as creating mutants of glycosyltransferases or altering glycosylation target sites through amino acid substitution, each come with limitations. Glycosyltransferase mutations may have pleiotropic effects on other proteins or pathways, even when genes are located within operons containing other flagella-related elements. Conversely, amino acid substitutions on flagellins may not fully recapitulate the structural or functional consequences of glycan removal and may affect flagellar integrity or function in unintended ways (23). Despite these constraints, these complementary strategies offer valuable insights and highlight the need for more refined tools to study glycosylation in situ. Future work should therefore focus on developing targeted approaches such as site-specific glycan editing or real-time imaging of motility and biofilm formation under flow conditions to better understand how glycosylation of surface structures like flagella and pili influences environmental survival and pathogen spread (56).

## Conclusion

Through laboratory column transport experiments, we investigated and quantified the transport and deposition behaviors of bacterial strains in sandy porous media, focusing on the role and glycosylation of flagella. By integrating observations with modeling, we elucidated the impact of flagella hydrophobicity on bacterial recovery and retention. Our findings clearly demonstrate that flagella impede bacterial transport and enhance deposition, with the WT showing higher retention compared to the Δ*fliC* strain, which exhibited greater potential for detachment. The Δ*fgt1* strain displayed faster transport and lower recovery rates, indicating that flagella glycosylation facilitates effluent recovery and reduces deposition. Furthermore, increased flagella hydrophobicity was found to hinder transport while promoting deposition. This study highlights critical insights into the effects of glycosylation, which are essential for evaluating the risk posed by phytopathogenic bacteria in farmland and for developing effective soil bioremediation strategies.

## Acknowledgment

The authors would like to acknowledge the gift of bacterial strains from Yuki Ichinose.

## Author Contributions

Conceptualization: Edvina Lamy, Yannick Rossez; Methodology: Edvina Lamy, Yannick Rossez; Formal analysis and investigation: Xin Zheng; Writing – original draft preparation: Xin Zheng; Writing – review and editing: Edvina Lamy, Yannick Rossez, Mounia Achak; Funding acquisition: Edvina Lamy, Yannick Rossez, Xin Zheng, Mounia Achak; Resources: Edvina Lamy, Yannick Rossez; Supervision: Edvina Lamy, Yannick Rossez.

## Ethics approval

Not applicable.

